# Hsp10 Targets the C-Terminus of α-Synuclein to Inhibit Secondary Nucleation and Remodel Labile Fibrils

**DOI:** 10.64898/2026.10.23.760823

**Authors:** Muhammed Shafeek Oliyantakath Hassan, Sydney Call, Ayobami Tosin Adegbenro, Jakob Villarreal, Marielle A. Wälti

## Abstract

Aggregation of α-synuclein is a central pathological feature of Parkinson’s disease and related synucleinopathies. The co-chaperonin Hsp10, a component of the Hsp60/Hsp10 chaperonin system, has recently been proposed to possess independent chaperone activity and has been found to co-localize with α-synuclein aggregates in diseased brains, suggesting a role in regulating α-synuclein assembly. Here, we combine solution NMR spectroscopy, cryo-electron microscopy, and kinetic analyses to define a multistage mechanism by which Hsp10 modulates α-synuclein aggregation. Hsp10 transiently interacts with monomeric α-synuclein, with residues on the inner surface and top-loop regions of Hsp10 primarily engaging the acidic C-terminal region of α-synuclein. Hsp10 exerts a concentration-dependent, biphasic effect on aggregation, accelerating fibrillization at substoichiometric concentrations while inhibiting de novo aggregation at higher concentrations through suppression of secondary nucleation. Furthermore, Hsp10 decorates fibril surfaces through interactions with the exposed α-synuclein C-terminus, thereby limiting secondary nucleation. Hsp10 also promotes the disaggregation of thermodynamically labile fibrillar fragments. Together, these findings establish a mechanistic framework for Hsp10-mediated regulation of α-synuclein assembly and expand the functional landscape of Hsp10 beyond its canonical role in mitochondrial protein folding.

## Introduction

The major pathological hallmark of Parkinson’s disease (PD) and related synucleinopathies is associated with the formation of intracellular inclusions known as Lewy bodies (LB) and Lewy neurites (LN) composed of abnormal aggregates of the protein alpha-synuclein (α-syn).^[1–2]^ These inclusions are particularly located in the *substantia nigra pars compacta* region of the brain and results in the degeneration of the dopaminergic neurons which intern results in characteristic motor symptoms of the disease including tremor, rigidity, bradykinesia, and postural instability.^[3–5]^ α-syn is an intrinsically disordered protein primarily expressed in the brain, particularly in the presynaptic terminals of the neuron and found in small amounts in other tissues.^[6]^ It is encoded by the SNCA gene and functions in the regulation of synaptic vesicle trafficking and the release of neurotransmitters.^[7]^ The protein consists of 140 amino acids and can be subdivided into three different domains. The N-terminal domain of α-syn, spanning residues 1–60, is highly positively charged and plays a crucial role in lipid binding.^[8–10]^ The central domain, known as the NAC (non-amyloid-β component) region (residues 61–95), is highly hydrophobic, forms the core of amyloid fibrils associated with various synucleinopathies, and plays a critical role in the aggregation process.^[10]^ The C-terminal domain of α-syn, containing residues 96-140, is highly acidic and influences fibril polymorphism.^[11–12]^ In the fibrillar state, the C-terminus and portions of the N-terminus form disordered and dynamic regions that extend outward from the fibril core, often referred to as the “fuzzy coat”.^[13]^ These surface-exposed intrinsically disordered regions serve as interaction sites for various cellular components, including molecular chaperones, and are important for α-syn transmission^[14]^.

Molecular chaperones are an ubiquitous and highly conserved class of endogenous proteins that are essential for maintaining protein homeostasis in cells of all organisms.^[15–16]^ They act as an essential cellular control mechanism, primarily by assisting in protein folding and by preventing the aggregation of improperly folded proteins.^[17–18]^ Heat shock proteins (Hsp) form the major class of molecular chaperones produced by cells in response to various stressful conditions.^[19–20]^ Among these, Hsp60 is a member of the chaperonin family that promotes protein folding within a central cavity, while Hsp10 functions as its co-chaperone by binding to Hsp60 and forming a lid over the folding chamber. ATP-dependent conformational cycling of the Hsp60/Hsp10 complex enables substrate encapsulation and provides a protected environment that can facilitate productive protein folding.^[21–23]^ Pathological studies have consistently demonstrated various Hsps, including Hsp60, to co-localize with α-syn aggregates in the characteristic LBs and LNs in the brain of synucleinopathy patients.^[24]^ Furthermore, functional studies have revealed that several Hsps effectively suppress α-syn aggregation.^[25–30]^ Additionally, specific chaperones demonstrate the ability to disaggregate the pre-formed fibrils.^[31–32]^

Recent studies suggest that Hsp10 possesses chaperone activity independent of its canonical association with Hsp60 and may directly influence the aggregation kinetics of α-syn.^[33–35]^ This possibility is particularly relevant to synucleinopathies, in which Hsp10 has been implicated in α-syn pathology. Szego et al. reported aberrant cytosolic accumulation of Hsp10 in synucleinopathies, with cytosolic trapping of mitochondrial Hsp10 representing an early pathological event. Proteomic analyses further identified Hsp10 as an α-syn-interacting protein in nerve terminals, while Hsp10 levels were reduced in synaptosomal fractions from patients with Parkinson’s disease, despite unchanged levels of α-syn and Hsp60; Hsp10 and α-syn were also found to colocalize.^[36]^ These observations implicate Hsp10 in α-syn homeostasis and raise the possibility that its mislocalization may alter α-syn assembly under pathological conditions. However, the molecular basis of the Hsp10–α-syn interaction and the mechanisms by which Hsp10 influences distinct stages of α-syn aggregation remain poorly understood.

In this study, we address this critical gap by systematically investigating the interaction between Hsp10 and α-syn at atomic resolution. Using solution NMR, cryo-EM, and other complementary biophysical techniques, we demonstrate that Hsp10 effectively inhibits de novo amyloid formation. Our data reveal that Hsp10 engages in a weak, transient interaction with the C-terminal domain of monomeric α-syn, thereby suppressing aggregation through inhibition of secondary nucleation. In addition, Hsp10 decorates α-syn fibrils in a highly ordered manner via interactions with their C-terminal regions. Hsp10 facilitates the disaggregation of sonicated, thermodynamically labile α-syn fibrils. Together, these findings provide a structural framework for understanding the interplay between Hsp10 and α-syn and expand the role of Hsp10 beyond mitochondrial proteostasis, identifying it as a key complementary component of the cellular defense machinery against synucleinopathies.

## Results and Discussion

### Hsp10 Inhibits α-synuclein Fibril Formation at High Concentration via Inhibition of Secondary Nucleation

Hsp10 has been shown to inhibit α-syn fibril formation.^[34]^ To verify these findings, we performed a Thioflavin T (ThT) assay. ThT is a benzothiazole dye that exhibits a marked increase in fluorescence upon binding to the repetitive β-sheet structures characteristic of amyloid fibrils, allowing fibril formation to be monitored over time. A total of 50 µM α-syn was incubated at 37 °C under shaking conditions in a buffer containing 10 mM sodium acetate at pH 5.0. Aggregation of α-syn began after approximately 7 hours, as indicated by an increase in fluorescence, and reached completion after 10–12 hours, with a half-time of fibril formation (T_1/2_) of 8.9 ± 1.5 hours, as determined by fitting using Amylofit^[37]^ (Figure 1a, blue). Negative stain electron micrographs indicate the formation of fibrils (Fig. 1b, blue). In contrast, in the presence of 50 µM or 150 µM Hsp10, the T_1/2_ increased to 9.9 ± 1.2 and 14.5 ± 0.71hours, respectively, indicating a clear inhibition of fibril formation (Figure 1a, cyan and red). On electron micrographs only few aggregates were visible in presence of 150 µM Hsp10. Since they were ThT negative, we conclude that these are non-fibrillar, amorphous aggregates (Fig. 1b, red). Notably, a buffer at pH 5 was used to accelerate aggregation. Hsp10 also inhibits α-syn aggregation at pH 7.4 in 10 mM Tris-HCl, 100 mM NaCl, and 1 mM EDTA (Figure S1). Surprisingly, at lower Hsp10 concentrations, aggregation was instead promoted: in the presence of 2 µM or 10 nM Hsp10, the T_1/2_ decreased to 7.4 ± 0.12 and 6.3 ± 0.28 hours, respectively (Figure 1a, green and orange). This enhancement of aggregation at sub-stoichiometric Hsp10 concentrations is consistent with previous observations reported for Hsp10-Aβ42 interactions, suggesting a concentration-dependent dual effect on fibril aggregation.^[33]^ Global kinetic analysis of the aggregation data was performed using AmyloFit, demonstrating that the compound selectively inhibits secondary nucleation.

**Figure 1.**
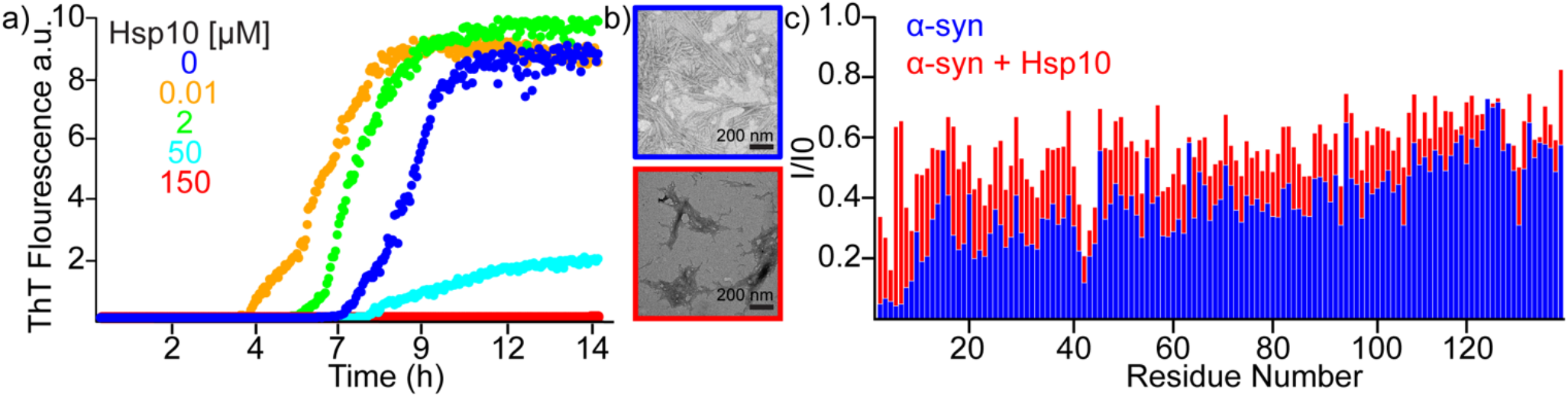
Hsp10 and α-syn interaction. a) ThT fluorescence assay of 50 μM α-syn monomer in the absence (blue) and presence of different concentrations of Hsp10, as indicated in the legend. Measurements were recorded at 37 °C under continuous shaking in 10 mM sodium acetate buffer (pH 5.0). b) Representative negative-stain TEM micrographs of α-syn after 15 h of incubation in the absence (blue) and presence of 150 μM Hsp10 (red). c) Normalized NMR signal intensity ratios (I/I_0_) obtained from a ^1^H,^15^N-HSQC spectrum of α-syn. I_0_ corresponds to the signal intensity of 80 μM monomeric ^15^N-labelled α-syn in the absence of fibril seeds, whereas I corresponds to the signal intensity of 80 μM monomeric ^15^N-labelled α-syn in the presence of 80 μM α-syn fibril seeds, in the absence (blue) or presence of 240 μM Hsp10 (red). Experiments were performed in 10 mM sodium phosphate buffer, 5% D_2_O, 0.03% NaN_3_, pH 6.5, at 298 K on an 800 MHz Bruker spectrometer.

To investigate how Hsp10 affects the secondary nucleation pathway of α-syn, we used ^1^H,^15^N-HSQC NMR spectroscopy. Unlabeled α-syn fibrils were first prepared by incubating 100 µM monomer in 3 mL fibrillization buffer at 37 °C with agitation at 800 rpm for one week. The resulting fibrils were collected by centrifugation (13,000 × g, 60 min, 25 °C), washed three times, and resuspended in 500 µL of 10 mM sodium phosphate buffer (pH 6.5). To generate fibril seeds, the fibrils were sonicated using a probe sonicator (Qsonica, Newtown, CT, USA) equipped with a microtip, using 10 cycles of 1 s on and 10 s off at 30% amplitude. The concentration of fibril seeds was determined by absorbance measurements at 280 nm following dissolution in 5 M guanidine hydrochloride. ^1^H,^15^N-HSQC experiments were then performed and analyzed using the following three samples: (i) 80 µM ^15^N-labeled α-syn monomer alone, which served as the reference spectrum (I_0_); (ii) 80 µM ^15^N-labeled α-syn in the presence of 80 µM unlabeled α-syn fibril seeds; and (iii) 80 µM ^15^N-labeled α-syn in the presence of 80 µM unlabeled fibril seeds and 240 µM unlabeled Hsp10. All spectra were recorded at 298 K on an 800 MHz Bruker spectrometer in 10 mM sodium phosphate buffer (pH 6.5) containing 5% D_2_O and 0.03% NaN_3_. Residue-specific normalized intensity ratios (I/I_0_) were calculated relative to the spectrum of ^15^N-labeled monomeric α-syn alone. Addition of α-syn fibril seeds to the monomeric α-syn resulted in substantial attenuation of HSQC peak intensities, particularly for residues within the N-terminal region of α-syn (Figure 1c, blue). This observation is consistent with a transient interaction between the N-terminal region of monomeric α-syn and the fibril surface, which has been implicated in secondary nucleation.^[38]^ In contrast, the presence of Hsp10 markedly reduced the seed-induced attenuation of the NMR signals (Figure 1c, red). This recovery of signal intensity suggests that Hsp10 limits the interaction of monomeric α-syn with the fibril surface, potentially through competition for overlapping binding sites, similar to mechanisms proposed for other chaperones, including BRICHOS.^[39]^ Together, these findings support a model in which Hsp10 associates with the fibril surface and reduces the accessibility of monomeric α-syn, thereby limiting secondary nucleation.

### Interaction of α-synuclein C-terminus with the Inner Surface of Bowl-Shaped Hsp10

To identify the binding site of Hsp10 on α-syn monomer, NMR-based chemical shift perturbation experiments were carried out using 80 μM ^15^N-labeled α-syn recorded in the absence and presence of 50–800 μM unlabelled Hsp10. The resulting perturbations were mapped onto the spectrum using BMRB accession numbers 18857 for α-syn and 53226 for Hsp10 assignments. Analysis revealed a clear binding site in the acidic C-terminal region, with the largest shifts observed at residues 123–140, and additional perturbations at residue 50 (Figure 2). Fitting of the titration curves using the software CcpNmr Analysis^[40]^ yielded a global dissociation constant (*K*_*D*_) of 472 ± 110 μM. Predominant binding to the C-terminus of α-syn was further confirmed using the C-terminally truncated α-syn (residues1-103) (CT-α-syn (1–103)). No chemical shift perturbations were observed upon addition of 100 μM unlabeled α-syn to 80 μM ^2^H,^15^N-labeled Hsp10 (Figure S2). Further chemical shift mapping from the perspective of Hsp10 was performed using 80 μM ^15^N-labeled and deuterated (^2^H,^15^N) Hsp10 in the absence and presence of 0–200 μM unlabelled α-syn (Figure 3). Significant perturbations were observed in residues 35, 36, 41–43, 53, 55, 58, 59, 63, 78, and 93, indicating a predominant binding at the inner surface of the bowl-shaped Hsp10 as well as at the top loop regions.

**Figure 2.**
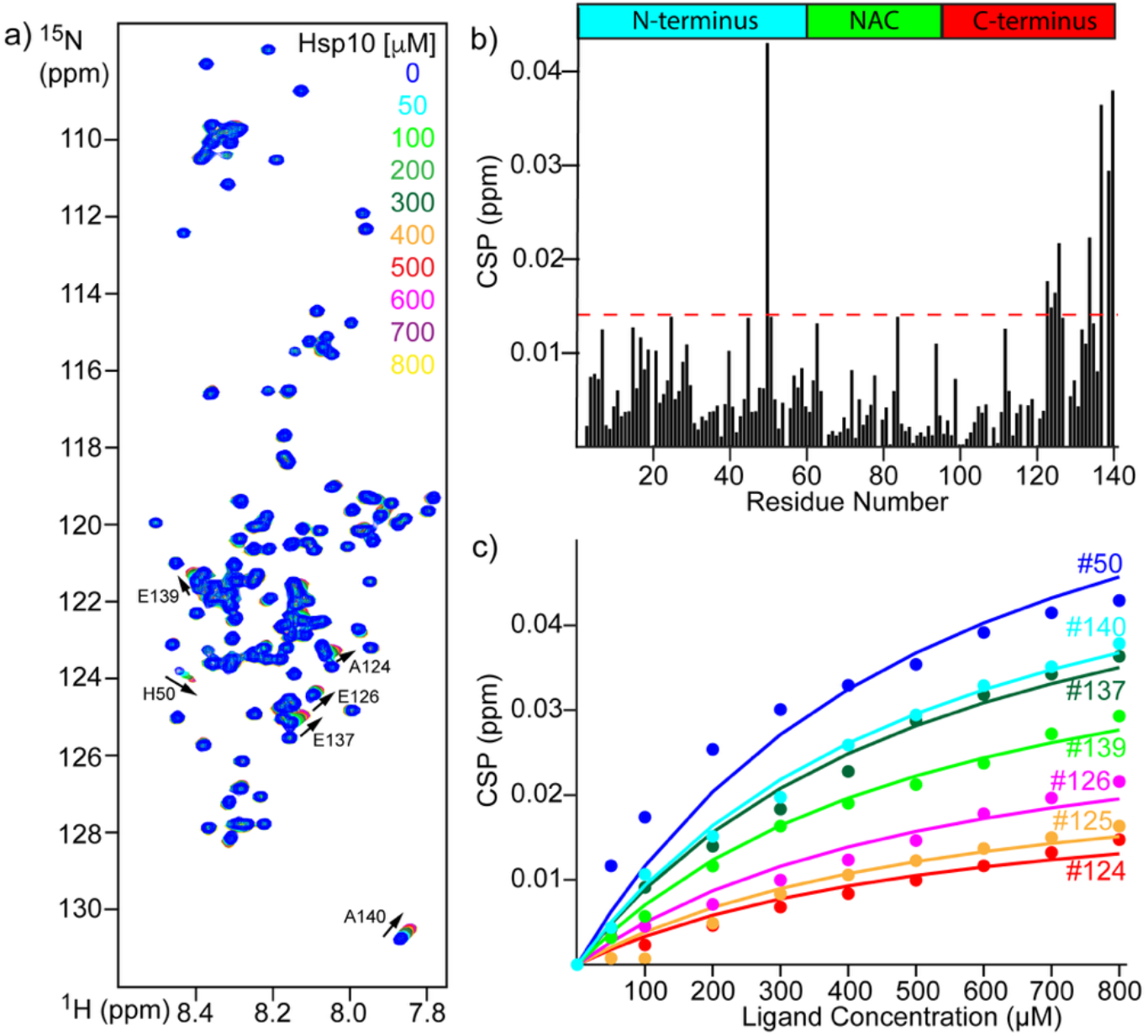
Interaction of Hsp10 with α-syn from the perspective of α-syn. a) NMR titration of 80 μM ^15^N-labeled α-syn in the presence of varying concentrations of Hsp10. b) Chemical shift perturbations of α-syn at the highest concentration of Hsp10, showing major shifts for the residues in the C-terminus and the residue at position 50. The dashed red line represents one standard deviation c) Fitting of the titration data. Residue numbers are indicated. The spectra were recorded at 800 MHz and 298 K in 10 mM sodium phosphate buffer containing 95% H_2_O, 5% D_2_O, and 0.03% NaN_3_ at pH 6.5.

**Figure 3.**
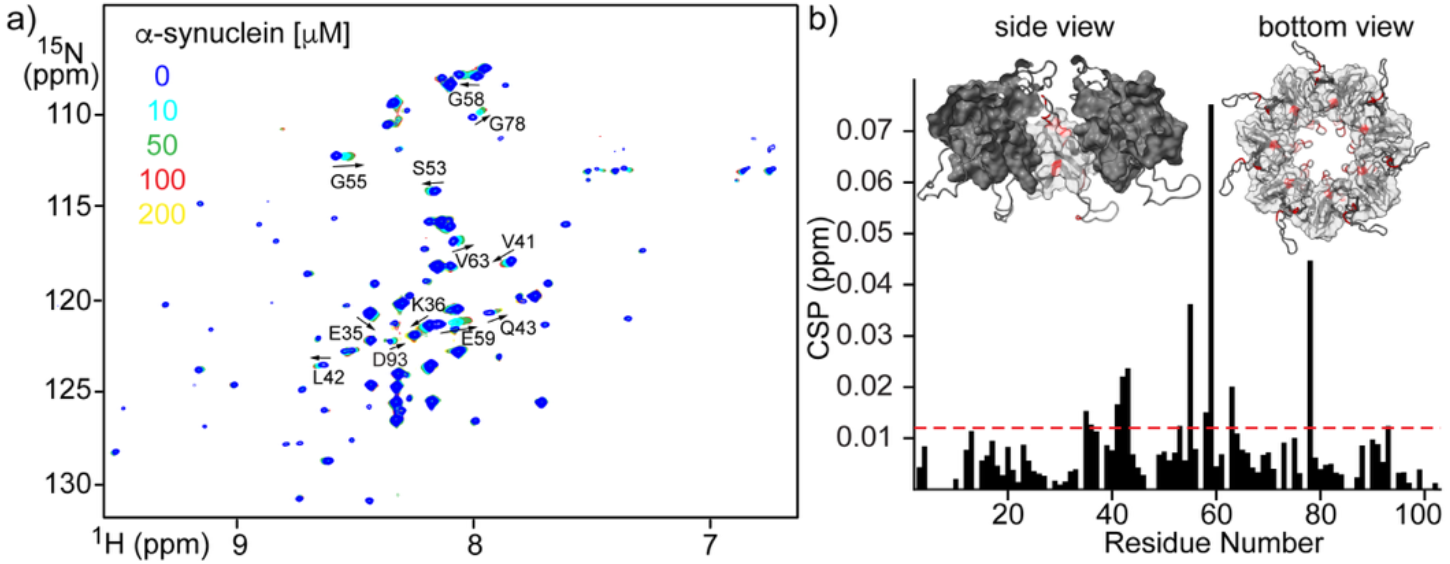
Interaction of Hsp10 with α-syn from the perspective of Hsp10. Chemical shift perturbations of 80 μM ^2^H,^15^N-labeled Hsp10 upon titration with 0–200 μM α-syn. a) Representative [^1^H,^15^N]-TROSY-HSQC spectra and b) the corresponding chemical shift perturbation profile. The red dashed line indicates one standard deviation above the mean perturbation, and residues exceeding this threshold are mapped onto the Hsp10 structure in the inset and highlighted in red. For clarity, one subunit was removed from the side view of the Hsp10 heptamer to reveal residues located within the central cavity. Experiments were recorded at 800 MHz and 298 K in 10 mM sodium phosphate buffer containing 95% H_2_O, 5% D_2_O, and 0.03% NaN_3_ at pH 6.5.

### α-Synuclein does not induce formation of single-ring Hsp10 species

Our previous data show that Hsp10 exists in equilibrium between single- and double-ring species.^[35]^ Previous reports by Larsson et al. have shown that Hsp10 can form stacked assemblies that were reported to disassemble into single-ring species upon addition of α-syn.^[34]^ Because solution NMR predominantly detects the single-ring species, this system is well suited to monitor structural transitions between different Hsp10 assemblies. To test whether α-syn induces the disassembly of Hsp10 assemblies, we compared signal intensities in ^1^H,^15^N TROSY-HSQC spectra of Hsp10 recorded in the absence and presence of α-syn. Upon addition of 200 µM unlabeled α-syn to 80 µM ^2^H,^15^N-labeled Hsp10, we observed a 36% decrease in NMR signal intensity (Figure S3a). Rather than indicating disassembly into smaller species, this decrease is consistent with the formation of larger or more slowly tumbling Hsp10 assemblies. Similarly, when 20 µM ^2^H,^15^N-labeled Hsp10 was mixed with 180 µM unlabeled α-syn, the signal intensity decreased by 12%, with no increase in signal intensity observed.

To further assess the oligomeric state of Hsp10, Blue Native PAGE experiments were performed. Samples containing 10 µM Hsp10 were mixed with 5, 10, 20, 40, or 50 µM α-syn and analyzed by Blue Native PAGE (Figure S3c). In the absence of α-syn, Hsp10 exhibited a single-ring-to-double-ring ratio of 5.4 ± 1.3. Upon addition of α-syn, this ratio decreased to 2.3 ± 0.4, 2.7 ± 1.0, 2.1 ± 0.2, 2.0 ± 0.3, and 2.2 ± 0.4, respectively (Figure S3d). Thus, the addition of α-syn at all concentrations tested resulted in a statistically significant decrease in the single-ring-to-double-ring ratio, consistent with an increased population of double-ring species. Together, the NMR and Blue Native PAGE data do not support a model in which α-syn induces the disassembly of Hsp10 into single-ring species. Instead, our results indicate that α-syn shifts the equilibrium toward double-ring Hsp10 assemblies.

### Hsp10 binds to and decorates α-synuclein fibrils

To investigate the interaction between Hsp10 and α-syn fibrils, SDS-PAGE analysis was performed using fibrils formed from both wild-type (wt) α-syn and C-terminally truncated α-syn (residues1-122) (CT-α-syn(1–122)) in the presence and absence of Hsp10. Fibrils were generated by incubating 100 µM α-syn in the absence or presence of 300 µM Hsp10 in fibrillization buffer (10 mM Tris, pH 7.4, 100 mM NaCl, 1 mM EDTA, and 0.03% NaN_3_) at 37 °C with shaking at 800 rpm for one week. Following fibril formation, samples were centrifuged at 13,000 × g for 60 min at 25 °C, and the resulting pellets were washed three times with fibrillization buffer to remove soluble proteins and non-specifically associated species. The washed pellets were subsequently denatured in fibrillization buffer containing 6 M urea, and both fibril samples and monomeric protein controls were adjusted to a final concentration of 5 µM prior to electrophoresis. Proteins were resolved by SDS-PAGE and visualized by Coomassie Blue staining.

SDS-PAGE analysis of the wt α-syn fibrillar pellet fraction revealed a clear Hsp10 band, demonstrating that Hsp10 remained associated with the fibrils following repeated washing steps (Figure 4a). Both the N- and C-terminal regions of α-syn are known to extend outward from the fibril core, forming a dynamic and solvent-accessible “fuzzy coat.” In contrast, Hsp10 was not detected in pellet fractions containing CT-α-syn(1–122) fibrils, indicating a loss of fibril association upon removal of the C-terminal region (Figure 4b). These findings suggest that Hsp10 interacts with α-syn fibrils through the C-terminal region, consistent with the binding mode observed for monomeric α-syn.

**Figure 4.**
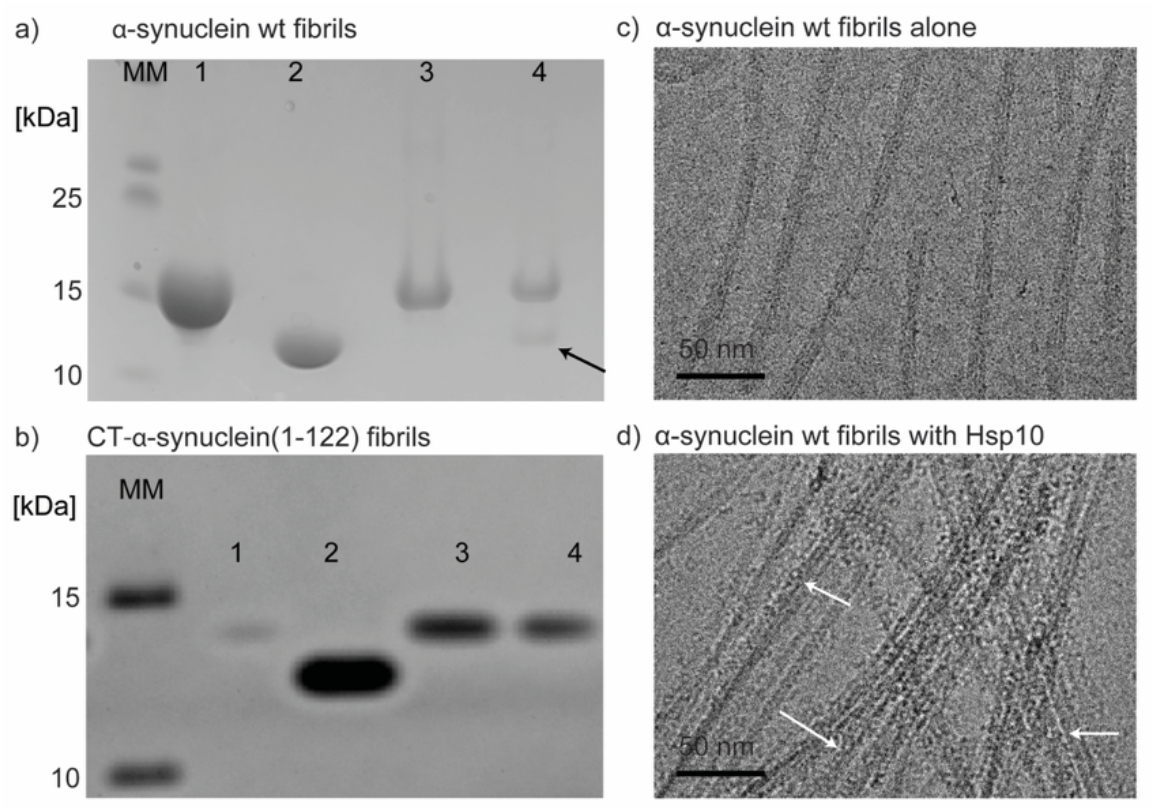
Hsp10 binding to α-syn fibrils. a, b) SDS–PAGE analysis of Hsp10 association with a) wild-type α-syn fibrils and b) CT-α-syn(1–122) fibrils. Fibrils were generated by incubating 100 µM α-syn in the absence or presence of 300 µM Hsp10. Prior to gel loading, all fibril samples were adjusted to a final α-syn concentration of 5 µM. MM, molecular weight marker; lane 1, α-syn; lane 2, Hsp10; lane 3, α-syn fibrils formed in the absence of Hsp10; lane 4, α-syn fibrils formed in the presence of Hsp10. The black arrow indicates Hsp10. c, d) Representative cryo-EM micrographs of 100 µM α-syn fibrils formed in c) the absence and d) presence of 300 µM Hsp10 in fibrillization buffer at 37 °C for one week with shaking at 800 rpm. White arrows indicate Hsp10 molecules bound to the fibril surface.

To directly visualize the interaction between Hsp10 and α-syn fibrils, cryo-electron microscopy (cryo-EM) was performed on fibrils prepared under identical conditions. Following fibrillization, samples were washed three times in 10 mM Tris, pH 7.4, 100 mM NaCl, 1 mM EDTA by repeated centrifugation and resuspension to remove soluble monomers and non-fibrillar aggregates. Subsequently, 5 µL of a 5 µM fibril sample was applied to a glow-discharged Quantifoil R1.2/1.3 Cu 200 mesh grid (Electron Microscopy Sciences). Grids were plunge-frozen in liquid ethane (−185 °C) using a Leica EM GP2 automated plunge freezer with a 10 s incubation time and a 2.5 s blot time. During grid screening, snapshots were collected, and the cryo-EM images revealed extensive and highly regular decoration of α-syn fibrils by Hsp10, with a ring-to-ring spacing of approximately 8 nm along the fibril axis (Figure 4d). This arrangement is reminiscent of the previously reported interaction between GroEL and HET-s fibrils.^[41]^ Consistent with the SDS-PAGE results, the cryo-EM micrographs demonstrated persistent association of Hsp10 with mature α-syn fibrils, indicating that Hsp10 binds stably to the fibril surface rather than interacting transiently with fibrillar species.

### Hsp10 promotes the disassembly of sonicated thermodynamically labile α-synuclein fibrils

Having established that Hsp10 binds mature α-syn fibrils, we next investigated whether Hsp10 can promote fibril disassembly. Because α-syn is known to adopt distinct fibril polymorphs depending on the aggregation conditions, we selected a fibril preparation that has previously been shown to be thermodynamically labile and, therefore, more amenable to disaggregation.^[42]^ Specifically, 150 μM monomeric α-syn was incubated under F65 polymorph-forming conditions (20 mM MES, 150 mM NaCl, 1 mM EDTA, pH 6.5) at 37 °C for 3 days with shaking at 800 rpm to induce fibril formation. The resulting fibrils were subsequently pelleted by centrifugation (13,000 × *g*, 60 min, 25 °C) and resuspended in an equal volume of disaggregation buffer (50 mM HEPES, pH 7.5, 50 mM KCl, 5 mM MgCl_2_, 2 mM DTT, and 2 mM ATP). The fibrils were then sonicated on ice in a 4 °C cold room using a Qsonica Q500 probe sonicator (six cycles of 10 s on and 30 s off; 30% amplitude) to generate smaller fibrillar seeds. The sonicated fibrils were diluted to a final concentration of 2.5 μM α-syn (monomer equivalent), and disassembly was monitored using a ThT fluorescence assay in the presence or absence of 25 μM Hsp10. ThT fluorescence was recorded every 5 min at 30 °C in 96-well plates using a BioTek Synergy Neo2 plate reader (excitation: 440 ± 10 nm; emission: 480 ± 10 nm). Each measurement cycle was preceded by 3 s of orbital shaking to maintain sample homogeneity (Figure 5a). The kinetic traces showed no significant change in ThT fluorescence over the 10 h observation period for sonicated α-syn fibrils incubated in the absence of Hsp10, indicating that the sonicated fibrils remained stable under these conditions. In contrast, the addition of Hsp10 resulted in a progressive decrease in ThT fluorescence, reaching 39.99 ± 0.02% of the initial fluorescence intensity after 2 h, after which the signal stabilized. This reduction in ThT fluorescence is consistent with a loss of β-sheet-rich fibrillar material and suggests that Hsp10 promotes the partial disassembly or structural remodeling of smaller, thermodynamically labile α-syn fibrillar species. In contrast, Hsp10 did not promote the disaggregation of mature α-syn fibrils. When fibrils were grown under the same conditions described above but were not sonicated, the addition of Hsp10 did not result in a change in ThT fluorescence intensity (Figure 5b). These results indicate that fragmentation of the fibrils into smaller fibrillar species by sonication is critical for Hsp10-mediated disassembly.

**Figure 5.**
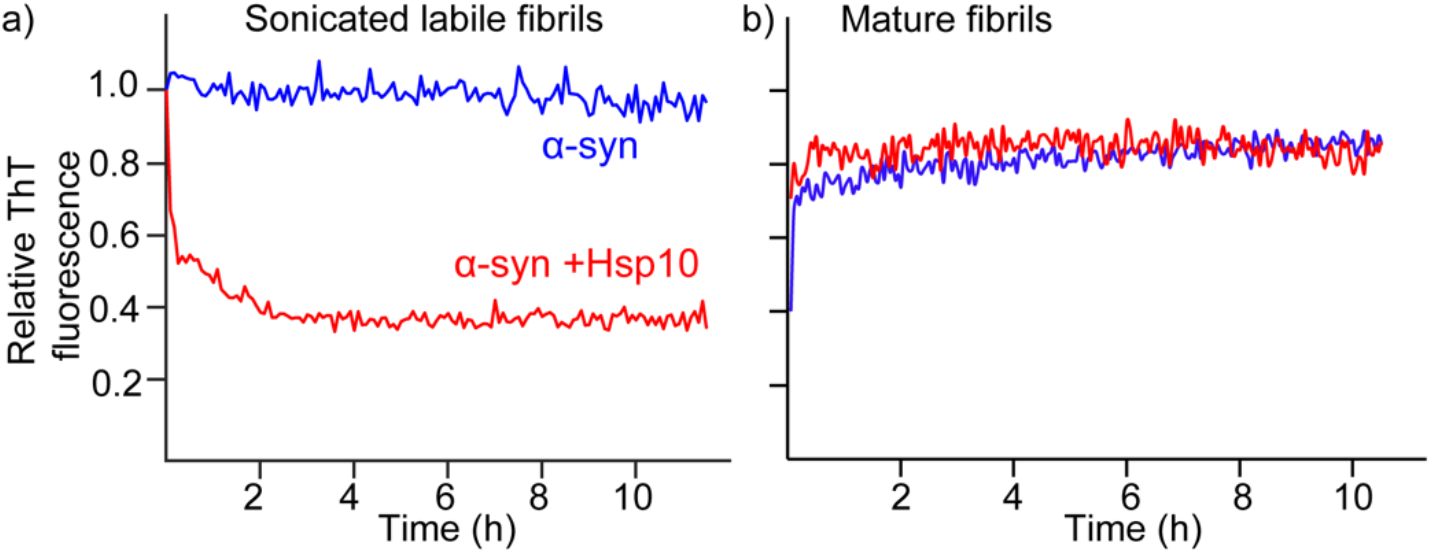
Disaggregation of α-syn fibrils. a) Normalized ThT fluorescence intensity over time for sonicated α-syn fibril seeds (2.5 μM, monomer equivalent) in disaggregation buffer (50 mM HEPES, pH 7.5, 50 mM KCl, 5 mM MgCl_2_, 2 mM DTT, and 2 mM ATP) containing 20 μM ThT. Blue curves represent fibril seeds alone, whereas red curves represent samples supplemented with 25 μM Hsp10. Fibrils were prepared under F65 polymorph-forming conditions (20 mM MES, 150 mM NaCl, 1 mM EDTA, pH 6.5) prior to sonication. b) ThT fluorescence intensity over time for mature, unsonicated α-syn fibrils (2.5 µM, monomer equivalent) in 10 mM Tris-HCl, 100 mM NaCl, and 1 mM EDTA at pH 7.4, containing 20 µM ThT, in the absence or presence of 25 µM Hsp10. The addition of Hsp10 did not result in a significant change in ThT fluorescence. Each data point represents the mean of three replicates.

## Conclusion

The reported association of Hsp10 with α-syn in nerve terminals, together with its altered distribution in synucleinopathies, suggests that Hsp10 may influence α-syn homeostasis beyond its canonical role within the Hsp60 chaperonin system.^[36]^ These observations provided the impetus to investigate whether Hsp10 directly engages α-syn and how such interactions affect the structural and kinetic transitions underlying its assembly. Our findings now establish a mechanistic framework for these interactions, revealing that Hsp10 can directly regulate α-syn across multiple stages of its assembly pathway.

In the present study, we establish Hsp10 as a direct and multistage regulator of α-syn aggregation and provide a mechanistic framework for how this small co-chaperonin modulates multiple stages of amyloid assembly. Our NMR data demonstrate that Hsp10 interacts predominantly with residues within the highly acidic C-terminal region of monomeric α-syn. The measured dissociation constant (*K*_*D*_ = 472 ± 110 μM) indicates relatively weak affinity, consistent with a transient interaction. Such weak interactions are characteristic of many ATP-independent holdase chaperones and may be advantageous for regulating intrinsically disordered proteins such as α-syn without permanently sequestering the monomeric state.^[28, 43–46]^ Our structural data further refine the molecular model of the Hsp10–α-syn interaction. Previous studies proposed an interaction between Hsp10 and the NAC region of α-syn; however, our NMR analyses indicate that the primary interaction site on monomeric α-syn is instead located within its C-terminal region. Moreover, our structural analyses show that α-syn interacts with the inner surface of the Hsp10 ring rather than predominantly with the exposed loop regions proposed previously.^[34]^ These findings raise the possibility that α-syn may access the Hsp10 inner cavity while leaving the Hsp10 loops available for Hsp60 binding. Together, these observations provide a structural framework for understanding how Hsp10 interacts with α-syn as an isolated co-chaperone.

A particularly striking finding of this study is the concentration-dependent, biphasic effect of Hsp10 on α-syn aggregation. Our ThT-based kinetic analyses revealed that sub-stoichiometric concentrations of Hsp10 accelerate fibrillization, whereas higher concentrations suppress α-syn aggregation. Similar concentration-dependent effects have previously been reported for Hsp10 in the aggregation of Aβ42 and the A30P variant of α-syn, suggesting that this dual behavior may represent a more general feature of Hsp10-mediated modulation of amyloid assembly. ^[33–34]^ Consistent with these observations, another study showed that Hsp10 promotes monomeric α-syn aggregation at low concentrations while inhibiting α-syn seeded aggregation.^[36]^ We hypothesize that the concentration-dependent effects of Hsp10 on α-syn aggregation arise, at least in part, from its interaction with the C-terminal region of α-syn. Monomeric α-syn exists as a heterogeneous conformational ensemble stabilized, in part, by transient long-range interactions between the acidic C-terminal region and the N-terminal and NAC regions, as demonstrated by NMR and molecular dynamics studies.^[47–52]^ Bertoncini et al. proposed that disruption of these long-range contacts can expose the NAC region and promote fibril formation. Consistent with a protective role for the C-terminus, C-terminal truncation or neutralization of the negatively charged C-terminal region accelerates α-syn aggregation relative to the full-length protein.^[11, 53–54]^ These findings support an important role for the C-terminal region in maintaining α-syn solubility and suppressing fibril formation. In this context, binding of Hsp10 to the C-terminal region could perturb the intramolecular interactions that stabilize the monomeric conformational ensemble and shift the equilibrium toward more open conformations. Such conformational rearrangements could increase the accessibility of the NAC region, thereby facilitating intermolecular interactions and promoting nucleation and fibril formation. This model provides a potential mechanistic explanation for the accelerated aggregation observed at low Hsp10 concentrations. In contrast, at higher Hsp10 concentrations, our kinetic analyses indicate that inhibition of secondary nucleation constitutes a major mechanism underlying the anti-aggregation activity of Hsp10. Secondary nucleation is a critical amplification pathway in amyloid assembly, as fibril surfaces can catalyze the formation of new oligomeric species that have been implicated as particularly toxic forms of α-syn.^[55–57]^ Cryo-EM revealed extensive and regular decoration of α-syn fibrils by Hsp10, supporting a model in which Hsp10 occupies fibril surfaces and interferes with the productive recruitment of soluble α-syn monomers. Because amyloid fibrils are composed of highly repetitive structures, they can present multiple repeated binding sites along their surface, allowing numerous Hsp10 molecules to associate with a single fibril. The regular arrangement of Hsp10 along the fibril axis further suggests an ordered interaction with exposed C-terminal regions within the fibrillar “fuzzy coat,” a region implicated in the cell-to-cell transmission and pathological propagation of α-syn assemblies.^[14]^ Similar fibril-surface interactions have been reported for other molecular chaperones and amyloid-binding proteins, including Hsp70 with α-syn fibrils, αB-crystallin, BRICHOS, Hsp104 with Aβ fibrils, and GroEL with HET-s fibrils.^[31, 41, 58–61]^ These observations suggest that direct binding and shielding of fibril surfaces may represent a broadly shared mechanism by which chaperones suppress amyloid amplification. Our NMR analyses provide additional mechanistic support for this model. In the presence of α-syn fibril seeds, the strongest reduction in monomer signal intensities occurred within the N-terminal region, consistent with transient interactions between soluble α-syn and the fibril surface during secondary nucleation. Notably, Hsp10 substantially reduced this signal attenuation, indicating that Hsp10 interferes with the interaction of soluble α-syn monomers with fibril surfaces. Together with the cryo-EM observations, these findings support a model in which Hsp10 suppresses secondary nucleation by occupying fibril surfaces and limiting the productive recruitment of α-syn monomers required for the generation of new aggregates. The inability of Hsp10 to bind fibrils generated from CT-α-syn(1–122) further supports the conclusion that the C-terminal region serves as the principal recognition site not only in monomeric α-syn but also within mature fibrillar assemblies.^[39]^

In addition to suppressing fibril formation, Hsp10 exhibited the ability to disaggregate pre-existing small fibrillar α-syn assemblies. ThT-based assays performed using α-syn fibrils generated under conditions known to produce thermodynamically labile fibrils, followed by sonication, revealed a decrease in fibril-associated fluorescence over time.^[42]^ These findings indicate that Hsp10 can also influence the stability or structural organization of aggregated α-syn species, although this activity appears to depend on the structural and thermodynamic properties of the fibrils.

Our findings also provide insight into the broader biological significance of Hsp10 in Parkinson’s disease and related synucleinopathies. Hsp10 has traditionally been characterized as the co-chaperonin partner of Hsp60 within the mitochondrial chaperonin system.^[21–23]^ However, increasing evidence indicates that Hsp10 can also exert chaperone-like functions independently of Hsp60.^[33–35]^ Our demonstration that Hsp10 directly interacts with both soluble and fibrillar α-syn suggests that these Hsp60-independent activities may extend to the regulation of α-syn assembly under pathological conditions. Given the central role of α-syn aggregation in synucleinopathies, Hsp10 may therefore represent an additional component of the cellular proteostasis network capable of modulating α-syn assembly.

In summary, our findings identify Hsp10 as an underappreciated regulator of α-syn assembly that acts through dynamic interactions with the exposed C-terminal regions of both monomeric and fibrillar species (Figure 6). Through this multistage mechanism, Hsp10 modulates fibril nucleation, suppresses secondary nucleation, interferes with fibril elongation, and influences the structural organization and stability of pre-formed aggregated species. Collectively, these findings support a model in which Hsp10 exerts stage-dependent effects on α-syn aggregation through concentration-dependent transitions between nucleation-promoting and fibril-shielding activities. These results expand the functional landscape of Hsp10 beyond its canonical role as a mitochondrial co-chaperonin and suggest a broader role for Hsp10 in regulating pathological protein aggregation associated with neurodegenerative disease.

**Figure 6.**
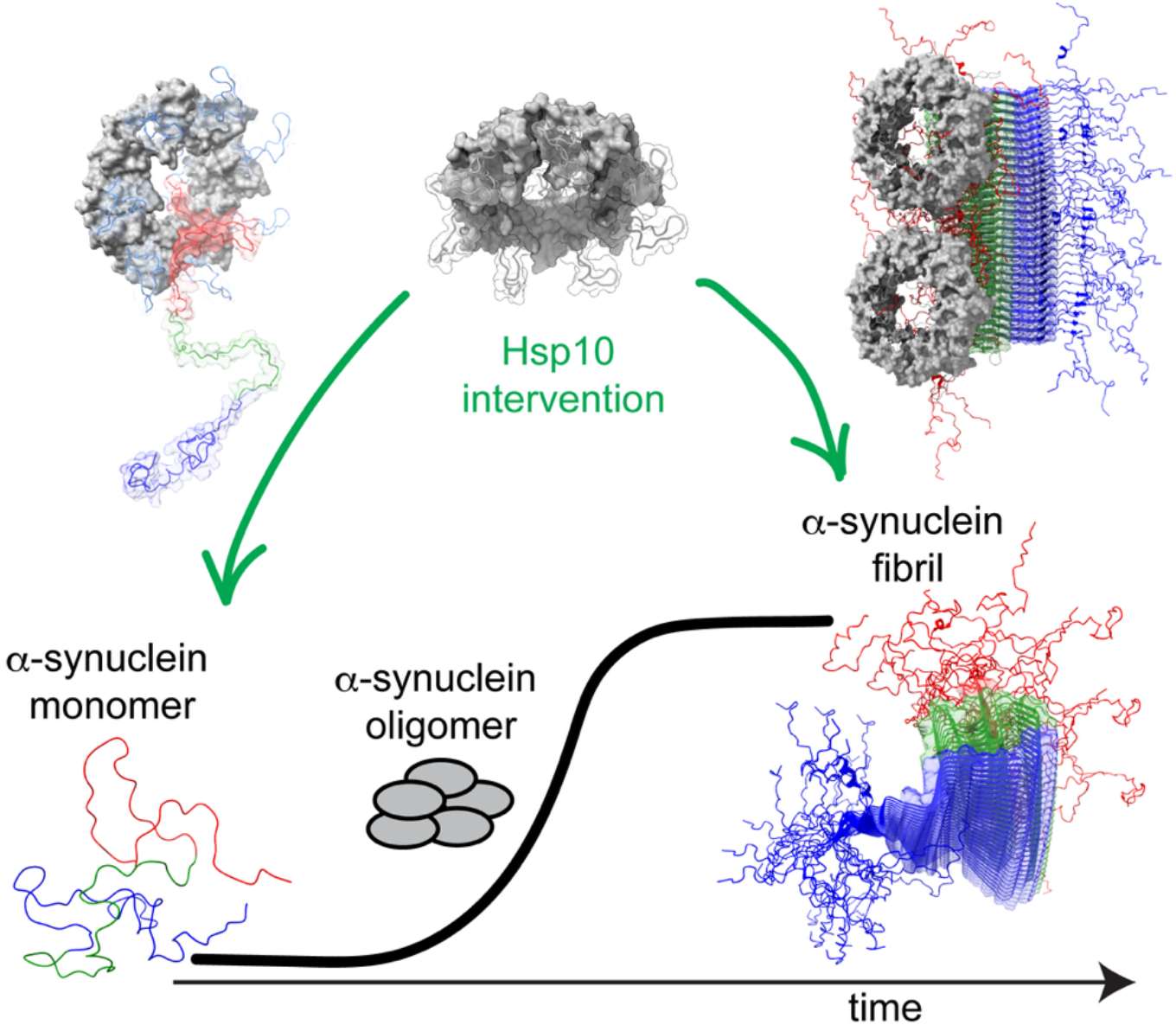
Summary of the α-syn–Hsp10 interaction. Hsp10 binds the C-terminal region of monomeric α-syn, with the interaction occurring at the inner face of the bowl-shaped Hsp10 oligomer, suggesting that α-syn could potentially be encapsulated within the Hsp60 cavity. Hsp10 also interacts with the C-terminal region of α-syn fibrils and inhibits secondary nucleation.

## Acknowledgement

This research was supported by the National Institute of General Medical Sciences of the National Institutes of Health under award number R00GM132496 and R35GM156786. The NMR was recorded at the University of Arizona - CBC Molecular Structure Core (MSC) NMR Facility, RRID: SCR_022888. A portion of this research was supported by NIH grant R24GM154185 and performed at the Pacific Northwest Center for Cryo-EM (PNCC) with assistance from Vamsee Rayaprolu. TEM measurements were performed at the John Cowley Center for High Resolution Transmission Electron Microscopy at Arizona State University with the assistance of Dwight Williams.

## Materials and Methods

### Expression and Purification of a-synuclein

Wild-type α-syn (Addgene plasmid #36046) was expressed in *E. coli* BL21 (DE3) competent cells. Cells were grown in LB medium supplemented with 100 μg/mL ampicillin at 37 °C. When the OD_600_ reached 1.2, the cultures were harvested by centrifugation (3,500 × *g*, 20 min, 25 °C), and the resulting cell pellets were gently resuspended in standard M9 minimal medium (containing D-(+)-glucose and NH_4_Cl). For isotopically labeled samples, M9 minimal medium was supplemented with the appropriate isotopic precursors (^15^N-NH_4_Cl and/or ^13^C-glucose; Cambridge Isotope Laboratories, Inc.). The cultures were allowed to recover for 1 h at 37 °C before induction with 1 mM IPTG and were incubated for an additional 8 h at 37 °C. Following expression, the cells were harvested by centrifugation (3,500 × *g*, 30 min, 25 °C). The purification protocol was adapted from Al-Azzani et al. with minor modifications.^[62]^ For cell lysis, cell pellets were resuspended in 50 mL of 10 mM Tris-HCl (pH 8.0), 1 mM EDTA, and 2 mM DTT. Streptomycin sulfate and protease inhibitors were added, and the suspension was stirred continuously at 4 °C for 20 min. Cells were then disrupted by three passes through an Avestin EmulsiFlex-C3 high-pressure homogenizer. The lysate was centrifuged at 23,500 × *g* for 10 min at 25 °C. The resulting supernatant was heated in a water bath at 95 °C for 15 min to remove heat-sensitive impurities, cooled to room temperature, and centrifuged at 23,500 × *g* for 15 min at 25 °C. The resulting supernatant was either frozen at −80 °C or immediately purified by anion-exchange chromatography using a HiTrap Q FF column (Cytiva). The column was equilibrated and washed with 50 mL of buffer A (10 mM Tris-HCl, 1 mM EDTA, 2 mM DTT, pH 8.0), and proteins were eluted using a 0–100% linear gradient of buffer B (10 mM Tris-HCl, 1 mM EDTA, 2 mM DTT, 1 M NaCl, pH 8.0) over a total volume of 50 mL. α-syn eluted at approximately 350 mM NaCl. Fractions containing α-syn were identified by sodium dodecyl sulfate–polyacrylamide gel electrophoresis (SDS–PAGE) and further purified by size-exclusion chromatography using a Superdex 75 Increase 10/300 GL column (Cytiva). The protein was eluted in 10 mM sodium phosphate buffer (pH 7.4), and fractions were analyzed by SDS–PAGE to confirm purity. Fractions containing monomeric α-syn (molecular weight, 14.4 kDa) were pooled, concentrated to approximately 500 μM using a 10 kDa molecular weight cutoff (MWCO) centrifugal concentrator, and fractionated in 300 µL aliquots and stored at −20 °C until further use. Protein concentration was determined by measuring absorbance at 280 nm using a NanoDrop spectrophotometer (Thermo Fisher Scientific). Purification of CT-α-syn(1–103) (Addgene plasmid #240597) required a modified chromatography strategy because of its higher isoelectric point.^[12]^ Cell lysis and heat treatment were performed as described above, except that the cell pellets were resuspended in 20 mM MES and 1 mM EDTA (pH 6.0), supplemented with protease inhibitor tablets and streptomycin sulfate. Following lysis by three passes through an Avestin EmulsiFlex-C3 high-pressure homogenizer, the lysate was clarified, heat-treated, and centrifuged a second time as described above. The resulting supernatant was either stored at −80 °C or immediately subjected to cation-exchange chromatography using an SP High Performance column (Cytiva). The column was equilibrated and washed with 50 mL of buffer A (20 mM MES and 1 mM EDTA, pH 6.0). Bound proteins were eluted using a 0–100% linear gradient of buffer B (20 mM MES, 1 mM EDTA, and 1 M NaCl, pH 6.0) over 50 mL. Fractions containing CT-α-syn(1–103) were identified by SDS–PAGE and further purified by size-exclusion chromatography using a Superdex 75 Increase 10/300 GL column (Cytiva) equilibrated with 10 mM sodium phosphate buffer (pH 7.4). Fractions corresponding to monomeric α-syn were identified by SDS–PAGE, pooled, and stored at −20 °C until further use.

### Expression and Purification of Hsp10

The plasmid expressing Hsp10 consisting of an N-terminal hexa-His tag was transformed into *E. Coli* BL21 (DE3) cells. The cells were grown in LB media supplemented with 100 μg/mL ampicillin resistance at 37 °C. The expression was induced with 0.5 mM IPTG once the OD_600_ of 0.8 was reached and incubated overnight at 20 °C. For the preparation of isotopically labelled samples, the cell culture in LB was centrifuged (3,500 x g, 20 minute, 25 ºC) after the OD_600_ reached 0.8, and the resulting cell pellets were resuspended in standard M9 minimal medium prepared in deuterium oxide, and supplemented with appropriate isotopic precursors (^15^N-NH_4_Cl and/or ^13^C-glucose, Cambridge isotopic laboratories Ltd.). The cell culture was allowed to recover for 1 hour at 37 °C before induction with 0.5 mM IPTG and incubated overnight at 20 °C. Following expression, the cells were harvested via centrifugation (3,500 xg for 30 minutes, 25 ºC) and resuspended to the lysis buffer (50 mM Tris, 300 mM NaCl, 10 mM Imidazole, pH 8) treated with streptomycin sulfate and protease inhibitor by continuous stirring at 4 °C for 20 minutes. The cells were disrupted by three subsequent passages through an Avestin EmulsiFlex-C3 high-pressure homogenizer, and centrifuged at 23,500 xg for 10 minutes at 25 ºC. The cell lysate was first subjected to nickel affinity chromatography (NTA-agarose, Cytiva). The target protein was eluted using a 50 mM Tris, 300 mM NaCl, 1 M Imidazole, pH 8. The fractions containing Hsp10 were dialyzed against an imidazole-free buffer (50 mM Tris, 100 mM NaCl, 10 mM Imidazole, 1 mM EDTA, 5 mM BME, 10% Glycerol, pH 8) and incubated with TEV protease overnight at 25 °C. Following cleavage, the sample was passed through a second Ni-NTA column. In this step, the cleaved target protein was collected in the flow-through. This fraction was further purified by size exclusion chromatography using a Superdex 75 Increase 10/300 GL column (Cytiva) in 50 mM sodium phosphate and 25 mM NaCl at pH 7.0. The protein concentration was measured with Absorbance at 280 nm and the purity confirmed using SDS-page. The final Hsp10 was aliquoted in 500 µL and stored at −20 °C at approximately 600 µM.

### Thioflavin-T Assay

Hsp10 and α-syn aliquots were removed from the freezer and thawed. Each sample was buffer-exchanged using a 3.5 kDa molecular weight cutoff dialysis cassette (Thermo Scientific) placed in a 1 L beaker containing the appropriate buffer: either 10 mM sodium acetate, pH 5.0, or 10 mM Tris-HCl, 1 mM EDTA, and 100 mM NaCl, pH 7.4. Samples were stirred at room temperature for at least 2 h before removal from the dialysis cassette, and protein concentrations were subsequently remeasured. Thioflavin T (ThT) was first dissolved in the appropriate buffer and filtered. The concentration of the filtered ThT solution was determined by measuring its absorbance at 412 nm, after which it was diluted to a final stock concentration of 700 µM. Samples were prepared containing 20 µM ThT, 50 µM α-syn, varying concentrations of Hsp10, and 0.03% NaN_3_. The samples were transferred to a 96-well plate (Corning 3651) and incubated in a BioTek Synergy Neo2 multimode plate reader. ThT fluorescence was monitored for 16 h at 37 °C with continuous orbital shaking. Fluorescence measurements were collected using an excitation wavelength of 450 nm and an emission wavelength of 482 nm.

#### NMR spectroscopy

All NMR experiments were conducted on Bruker Avance Neo spectrometers (600 MHz and 800 MHz), equipped with TCI z-axis gradient cryoprobes, at a temperature of 298 K. NMR data was processed with NMRPipe^[63]^ and analyzed using CcpNMR.^[40]^

### Chemical Shift Perturbation Mapping Experiments

Chemical shift perturbation (CSP) experiments were carried out in the NMR buffer (10 mM sodium phosphate, pH 6.5, in 95% H_2_O, 5% D_2_O). To minimize pH-dependent chemical shift variations during titration, both α-syn and Hsp10 samples were removed from the freezer, thawed, and buffer-exchanged. Each sample was placed in a 3.5 kDa dialysis cassette (Thermoscientific), and placed into a 1 L beaker containing the NMR buffer. The samples were left to stir in the buffer for at least 2 hours at room temperature, before being removed. The concentration was determined by absorbance at 280 nm. To map the binding sites of Hsp10 on α-syn, unlabeled Hsp10 was titrated into 100 µM ^15^N-labeled α-syn at final concentrations ranging from 0 to 800 µM. Conversely, to map the binding sites of α-syn on Hsp10, unlabeled α-syn was titrated into 100 µM U-[^2^H,^15^N]-labeled Hsp10 at final concentrations ranging from 0 to 1 mM. For the titration involving ^15^N-labeled α-syn and unlabeled Hsp10, ^1^H/^15^N HSQC spectra were acquired at each titration point. For the titration involving U-[^2^H,^15^N]-labeled Hsp10 and unlabeled α-syn, [^1^H,^15^N] TROSY-HSQC spectra were collected at each corresponding titration point. The assignment of Hsp10 and α-syn were taken from BMRB: (53226 and 18857 respectively). The pH of all samples was monitored using a SevenDirect SD20 pH meter equipped with an InLab NMR pH sensor (Mettler Toledo, Columbus, OH, USA). All measurements were maintained within ±0.01 pH units.

### Chemical Shift Perturbation Calculation

Chemical shift perturbations (CSP) were calculated using the following equation:

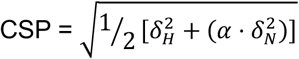

where *δ*_H_ and *δ*_N_ represent the changes in the proton and nitrogen chemical shifts, respectively, and α is the scaling factor for the nitrogen shift (*α* = 0.2).

### Dissociation Constant (*K*_D_) Calculation

The dissociation constant (*K*_*D*_) was determined by global fitting of the CSP data from all residues showing significant perturbation. The fitting was performed using the following equation:

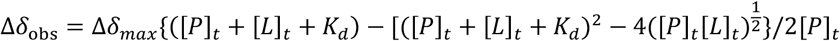

where Δ*δ*_obs_ is the observed change in chemical shift from the free state, Δ*δ*_max_ represents the maximum shift change upon saturation, and [*P*]_t_ and [*L*]_t_ are the total concentrations of the protein and ligand, respectively.

### Preparation of α-syn fibrils and seeds

Hsp10 and α-syn aliqouts were removed from the freezer and thawed. The samples were then buffer-exchanged by placing in a 3.5 kDa dialysis cassette (Thermoscientific), which was allowed to stir in 1 L of buffer (10 mM Tris, pH 7.4, 100 mM NaCl, 1 mM EDTA) for at least 2 hours at room temperature. The α-syn fibrils were prepared by incubating 100 µM α-syn with and without 300 µM Hsp10 in fibrillization buffer (10 mM Tris Buffer at pH 7.4, 100 mM NaCl, 1 mM EDTA, and 0.03% NaN_3_) at 37 °C for one week with shaking at 800 rpm. The fibrils were washed three times in the same buffer with resuspension and centrifugation at 13000 xg for 45 minutes to remove any soluble monomers and non-fibrillar aggregates.

To prepare thermodynamically labile fibril segments, 100 µM α-syn monomers were incubated in 20 mM MES, 150 mM NaCl, and 1 mM EDTA at pH 6.5.^[42]^ The samples were left to aggregate at 37 °C under continuous shaking at 800 rpm for 3 days. Following the aggregation period, the fibrillar fractions were isolated via centrifugation at 13,00 xg for 45 min at 25 °C. The resulting pellets were washed three times with the incubation buffer to remove non-fibrillar species and non-specifically bound proteins.

### Disaggregation

The fibrils are centrifuged at 13000 xg for 45 min at 25 °C. The resulting pellets are washed three times in 10 mM Tris-HCl, 100 mM NaCl, and 1 mM EDTA at pH 7.4. The resulting concentration was obtained by dissolving the pellet in 6 M urea and measuring the absorbance at 280 nm. Fibril seeds were generated using a Qsonica-Q500 ultrasonic probe sonicator (Qsonica, Newtown, CT, USA) (6 cycles of 10 s on, 30 s off, amplitude: 30%). The disaggregation reaction was monitored via a Thioflavin T fluorescence (ThT) assay in a total reaction volume of 100 μL. Reactions consisted of 2.5 μM fibril seeds (monomer equivalent) incubated with 25 μM Hsp10 and 20 µM ThT at 30 °C for 10 min For the mature, non-sonicated a-syn fibrils sample, the fibrils were centrifuged at 13,000 × g for 45 min at 25 °C, and the resulting pellets were washed three times and resuspended in 10 mM Tris-HCl, 100 mM NaCl, and 1 mM EDTA at pH 7.4. For the ThT disaggregation assay, α-syn fibrils were diluted to a final concentration of 2.5 µM (monomer equivalent) and incubated in the presence or absence of 25 µM Hsp10 and 20 µM ThT in a total reaction volume of 100 µL. All samples were loaded onto a 96-well assay plate (Corning 3651). The assays were conducted at 30°C for 12 h with the fluorescence intensity recorded using a BioTek Synergy Neo2 plate reader (Agilent Technologies, Santa Clara, CA, USA) with excitation and emission wavelengths set to 440-10 nm and 480-10 nm, respectively. Data points were aquired in 3 min intervals, with each cycle preceded by 3 s of orbital shaking to maintain sample suspension.

### Electron Microscopy

For Transmission Electron Microscopy, the fibrils were diluted in 10 mM Tris, pH 7.4, 100 mM NaCl, 1 mM EDTA to a monomeric concentration of 10 μM. 4 μL of the diluted sample was then blotted on a 400-mesh carbon-coated copper grid (Electron Microscopy Sciences) for 1 min, washed twice with deionized water, and stained with 10 μL of VitroEase Methylamine Tungstate Negative Stain (Electron Microscopy Sciences) for 1 min. The images were recorded using a TALOS L120C (TFS) transmission electron microscope operating at 120 kV. For Cryo-EM, the samples were kept at room temperature prior to freezing. 4 μL of sample was applied to a glow-discharged (PELCO easiglow) QF-R 1.2/1.3 Cu 200 M grid (Electron Microscopy Sciences). Grids were frozen into −185 °C liquid ethane using a Leica EM GP2 automatic plunge freezer equilibrated to 80% humidity and 10 °C with a 10 s sample incubation time and a 3 s blot time on Whatman 1 paper. Sample acquisition was conducted at the Pacific Northwest Center for Cryo-EM on a GLACIOS 2 200kV transmission electron microscope with a Thermo Scientific Falcon 4i Direct Electron Detector.

### Electrophoretic Analysis of Protein Samples

Immediately following purification, samples were analyzed by SDS-PAGE (4–12%) to assess protein purity. To determine changes in the single-ring-to-double-ring ratio of Hsp10 upon addition of α-syn, samples were analyzed by Native-PAGE (4–16%). Samples were thawed and diluted in 10 mM Tris, 150 mM NaCl, pH 7.4. For both SDS-PAGE and Native-PAGE, proteins were visualized by Coomassie Blue staining. For fibrillar samples, α-syn wild-type and C-terminally truncated α-syn (residues 1–122) fibrils were generated under F65 polymorph conditions. α-syn monomers (50 μM) were incubated in 20 mM MES buffer, pH 6.5, containing 150 mM NaCl and 1 mM EDTA, either in the absence or presence of Hsp10 at a 1:3 α-syn molar ratio in monomeric units. Following the aggregation period, fibrillar fractions were isolated by centrifugation at 13,000 × g for 60 min at 25 °C. The resulting pellets were washed three times with buffer to remove non-fibrillar aggregates and excess unbound Hsp10. To resolve the constituent protein components, the final washed pellets were denatured in 6 M urea to a final monomer-equivalent concentration of 5 μM, matching that of the monomeric controls, and analyzed by SDS-PAGE. The gel intensities were quantified using ImageJ.^[64]^

