## Supplementary Information for "Hsp10 Targets the C-Terminus of α-Synuclein to Inhibit Secondary Nucleation and Remodel Labile Fibrils"

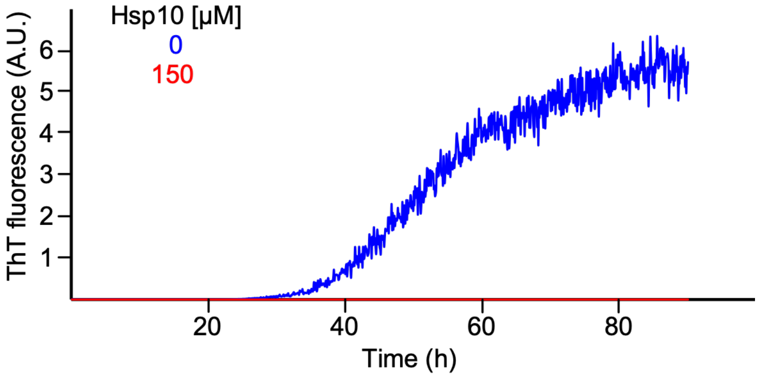


**Figure S1.** ThT fluorescence assay of 150 μM α-synuclein monomer in the absence (blue) and presence (red) of 150 μM Hsp10 over time, recorded at 37 °C under shaking conditions. The buffer contained 10 mM Tris, 100 mM NaCl, 1 mM EDTA, 0.03% $NaN_3$, and 20 μM ThT at pH 7.4.

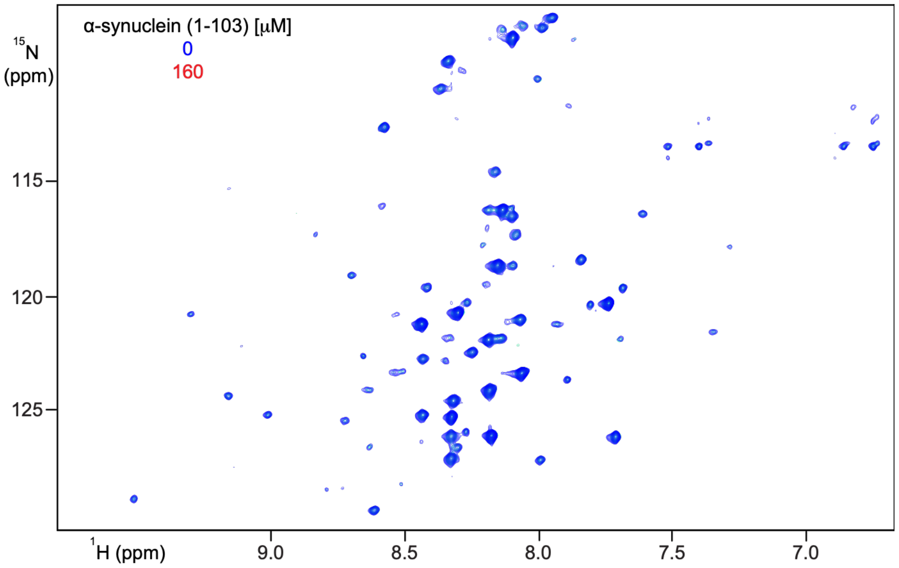


**Figure S2.** 2D [$^1$H, $^{15}$N]-TROSY HSQC NMR spectra of 80 μM $^2$H,$^{15}$N-labelled Hsp10 in the absence (blue) and presence (red) of 100 μM unlabeled C-terminally truncated (CT) α-syn (1-103). The experiment was recorded at 298 K in 10 mM $NaPO_4$, 5% $D_2O$, and 0.03% $NaN_3$ at pH 6.5 at 800 MHz.

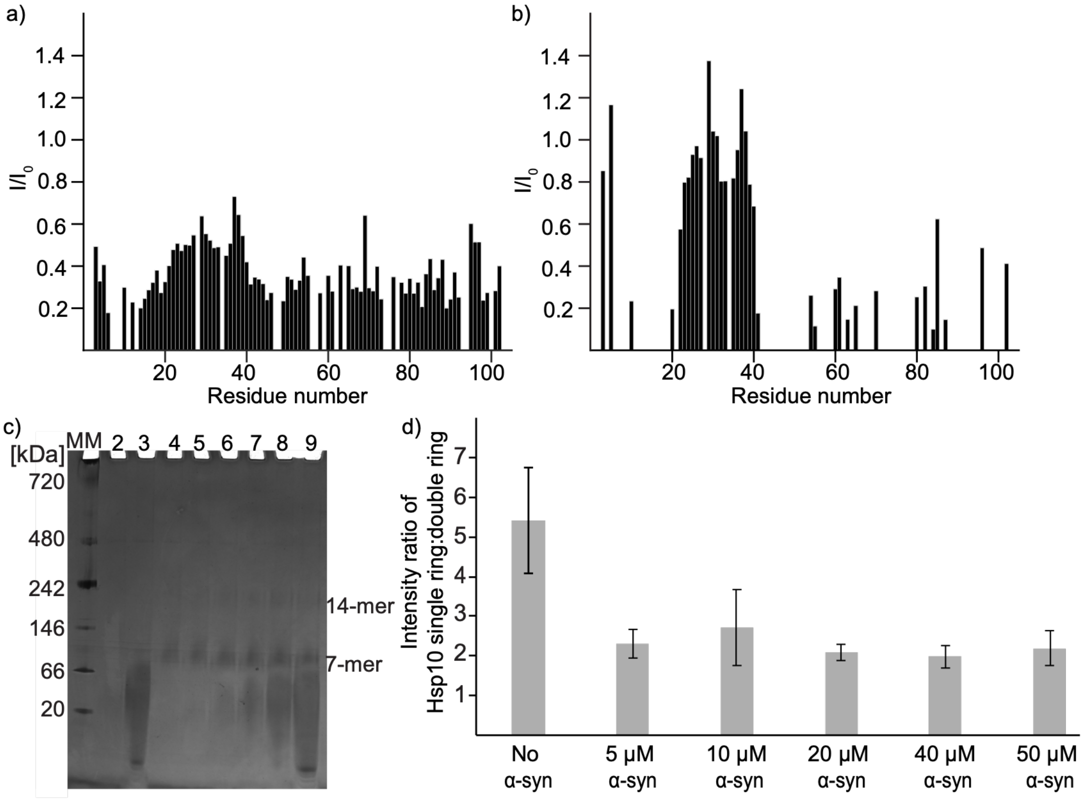


**Figure S3.** Lack of Hsp10 disassembly into single-ring in presence of α-syn. Intensity ratio of 80 μM $^2$H,$^{15}$N labeled Hsp10 in absence (*I₀*) and presence of 200 μM unlabeled α-syn (*I*) and b) 20 μM $^2$H,$^{15}$N labeled Hsp10 in absence (*I₀*) and presence of 180 μM unlabeled α-syn. The experiment was recorded at 298 K in 10 mM $NaPO_4$, 5% $D_2O$, and 0.03% $NaN_3$ at pH 6.5 at 800 MHz. c) Native-PAGE (4–16%) analysis of Hsp10 at varying α-synuclein concentrations. Lane 1: Molecular weight marker. Lane 2: 5 μM α-syn, Lane 3: 50 μM α-syn only, Lane 4: 10 μM Hsp10 only, Lane 5-9: 10 μM Hsp10 with 5, 10, 20, 40, and 50 μM α-syn. d) Quantification of the Hsp10 single-ring-to-double-ring intensity ratios derived from the gels shown in panel (c) using ImageJ.[1] Error bars represent the standard deviation.
